# Structural basis for substrate gripping and translocation by the ClpB AAA+ disaggregase

**DOI:** 10.1101/428458

**Authors:** Alexandrea N. Rizo, JiaBei Lin, Stephanie N. Gates, Eric Tse, Stephen M. Bart, Laura M. Castellano, Frank DiMaio, James Shorter, Daniel R. Southworth

**Affiliations:** Graduate Program in Chemical Biology, University of Michigan, Ann Arbor, MI, USA; Department of Biochemistry and Biophysics, Institute for Neurodegenerative Diseases, University of California, San Francisco, CA 94158, USA; Department of Biochemistry and Biophysics, Perelman School of Medicine at the University of Pennsylvania, Philadelphia, PA 19104, U.S.A.; Department of Biochemistry, University of Washington, Seattle, WA 98195, USA

## Abstract

Bacterial ClpB and yeast Hsp104 are homologous Hsp100 protein disaggregases that serve critical functions in proteostasis by solubilizing protein aggregates. Two AAA+ nucleotide binding domains (NBDs) power polypeptide translocation through a central channel comprised of a hexameric spiral of protomers that contact substrate via conserved pore-loop interactions. To elucidate the translocation mechanism, we determined the cryo-EM structure of a hyperactive ClpB variant to 2.9 Å resolution bound to the model substrate, casein in the presence of slowly hydrolysable ATPγS. Distinct substrate-gripping mechanisms are identified for NBD1 and NBD2 pore loops. A trimer of N-terminal domains define a channel entrance that binds the polypeptide substrate adjacent the topmost NBD1 contact. NBD conformations at the spiral seam reveal how ATP hydrolysis and substrate engagement or disengagement are precisely tuned to drive a stepwise translocation cycle.

## INTRODUCTION

Heat shock protein (Hsp) 100 protein complexes are critical cell stress responders that solubilize and unfold toxic protein aggregates and amyloids, thereby enhancing thermal and chemical tolerances^1,2^. They are members of the conserved family of AAA+ molecular machines that form dynamic hexameric-ring structures and undergo ATP hydrolysis-driven translocation of polypeptide substrates through a central channel^3–6^. Homologs ClpB from bacteria^7^, and Hsp104 from yeast^8^ are powered by two AAA+ nucleotide-binding domains (NBD1 and NBD2) per protomer and collaborate with the Hsp70 chaperone system to promote disaggregation and downstream re-folding of substrates^9,10^. Hsp104 recognizes and unfolds amyloid structures and is required for the transmission of yeast prions, such as Sup35, thereby enhancing prion propagation, and enabling selective advantages^11–13^. Pore-loop motifs within the AAA+ domains are essential for translocation and contain key Tyr residues that are a part of a highly conserved aromatic/hydrophobic pair proposed to interact and grip polypeptide substrates^14–17^. ATP hydrolysis requires conserved “Walker A” and “Walker B” motifs and an Arg finger residue in an adjacent protomer that contacts the nucleotide and contributes to interdomain communication^18^. How substrate interactions and ATP hydrolysis are coordinated across the domains and around the hexamer to drive polypeptide translocation remain ongoing questions for this conserved class of AAA+ machines.

An α-helical coiled-coil middle domain (MD) within the NBD1 of ClpB and Hsp104 wraps equatorially around the hexamer^8,19–21^ and interacts with Hsp70 to facilitate substrate loading^22,23^. The MD undergoes nucleotide-dependent conformational changes^6,20,24^ and is a key allosteric modulator of disaggregase function^25,26^. Point mutations in the MD alter ATP hydrolysis and potentiate activity^27^. Indeed, MD variants have been identified that enable Hsp104 to more effectively dissolve self-templating fibrils formed by human neurodegenerative disease proteins including α-synuclein, TDP43, FUS, and TAF15^25,28–30^. In addition, amino-terminal domains (NTDs) are connected to the NBD1 by a flexible linker^7^ and form an additional ring in the hexamer^31^ that may facilitate substrate transfer to the AAA+ domains. Substrate interaction sites have been identifed^32^ and the NTDs are critical for the dissolution of prions by Hsp104^33^, supporting plasticity in the mechanism, however, specific functions during translocation are unknown.

Recent cryo-EM structures of substrate-bound Hsp104^6^ and ClpB^5^ complexes identify an asymmetric architecture of the hexamer and a remarkable spiral of pore loop-substrate contacts by both NBD1 and NBD2. Five protomers contact an 80 Å-long unfolded polypeptide strand via the conserved Tyr residues with an approximate 6.5 Å, dipeptide spacing while a sixth protomer at the spiral seam is unbound to substrate and flexible^6^. Pore loop-substrate interactions are similarly spaced for both NBD1 and NBD2 rings in Hsp104^6^, but are different for ClpB, with NBD1 adopting a more planar arrangement compared to NBD2^5^. The architecture and substrate interactions are similar to other recent structures of single and double-ring AAA+ complexes^34–36^, supporting a conserved substrate interaction mechanism. However, for the double-ring translocases, higher resolution structures are needed to further define substrate interactions and identify how coordination between domains drives translocation. In the cryo-EM dataset of Hsp104 we identified an additional conformation in which the seam protomer rearranges to contact the substrate at the next position, resulting in a complete spiral of contacts by the hexamer, and, together, supports a two amino acid-step rotary translocation mechanism^6^. This extended state was not identified in ClpB. Additionally, substrate-free structures of Hsp104^6,20^ with ADP and the nonhydrolyzable ATP analog, AMPPNP, and of VAT^37^, an archeal AAA+ unfoldase, identify a dramatically different open “lock washer” conformation of the hexamer that may function during substrate engagement or release. While the Hsp104 structures were of wild type (WT), ClpB and other structures relied on Walker B mutants that are stabilized for substrate and nucleotide binding but inactive for hydrolysis and disaggregation^38^, leading to questions about which conformations represent active states that are required for substrate translocation.

To elucidate conserved mechanisms of disaggregation we sought to determine high-resolution cryo-EM structures of substrate-bound ClpB that identify pore loop-substrate interactions and conformational states involved in translocation. Utilizing the model substrate casein and the slowly hydrolysable ATP analog, ATPγS we previously established for stabilizing substrate-bound Hsp104^6^, we determined the structure of a hyperactive ClpB variant^27^, ClpB^K476C^, to 2.9 Å resolution. At this resolution, the spiral array of substrate contacts are precisely defined, revealing that the NBD1 and NBD2 have different substrate interaction “gripping” mechanisms that are coordinated by two distinct pore loop motifs. We identify that key charged residues in the NBD1 form a network of stabilizing cross-pore loop contacts and while secondary pore loops in both domains (D1’ and D2’) are identified to make additional substrate contacts and stabilizing interactions. Modeling of the well-resolved density for the casein substrate reveals hydrophobic sequence characteristics of the substrate that are preferred by the NBD1 and NBD2. By Focused classification and refinement we identify a trimer of N-terminal domains that form a substrate entrance channel and position the polypeptide above the NBD1 to facilitate delivery. The protomers at the spiral seam of the hexamer undergo conformational and nucleotide-state changes that are on-path with hydrolysis-driven, stepwise translocation of the polypeptide. Additionally, the NBD1-NBD2 interface undergoes an expansion at the spiral seam we propose to function as a coordinated mechanism for substrate pulling and release during the disaggregation cycle.

## RESULTS

### Cryo-EM structure of the hyperactive disaggregase complex: ATPγS-ClpB^K476C^:casein

Previous structures of ClpB and Hsp104 identified a right-handed spiral arrangement of NBD1 and NBD2 pore loops contacting the polypeptide substrate along the channel^5,6^, however higher resolution structures are needed to better define the contacts and substrate interaction mechanism. Additionally, previous structures contained Walker B mutations^5,34,35^ that bind substrate but are inactive for ATP hydrolysis. Indeed, the ClpB: casein structure^5^ utilized a “BAP” variant (where residues S722-N748 are replaced with V609-I635 of ClpA^3^) containing a Walker B mutation in both NBD1 and NBD2. Thus, structures of active complexes are needed to establish the translocation mechanism. Similar to previous studies, fluorescein-labeled (FITC) casein, a well-established unfolded model substrate, and ATPγS, a slowly hydrolysable analog that can power translocation of unfolded polypeptides *in vitro*^39,40^, were used for investigating active, substrate-bound complexes. WT ClpB was initially tested for substrate binding by size exclusion chromatography (SEC) (Supplementary Fig. 1a), however its stability was problematic under cryo-EM conditions and initial reconstructions did not go to high-resolution (data not shown). The 2D cryo-EM averages of the fractionated sample did show predominantly “closed”, substrate bound conformations (Supplementary Fig. 1b). To identify a more stable, but active complex, a hyperactive ClpB variant containing a K476C mutation in helix L2 of the MD was tested^27^. ClpB^K476C^ bound substrate similarly as WT by SEC, indicated by the overlapping elution peaks for FITC-casein and ClpB^K476C^ (Supplementary Fig. 1a). Reference-free class averages show well-defined features, different orientations and an architecture that is consistent with WT and the previously characterized substrate bound “closed” conformation of Hsp104^6^ (Supplementary Fig. 1b).

**Fig. 1:**
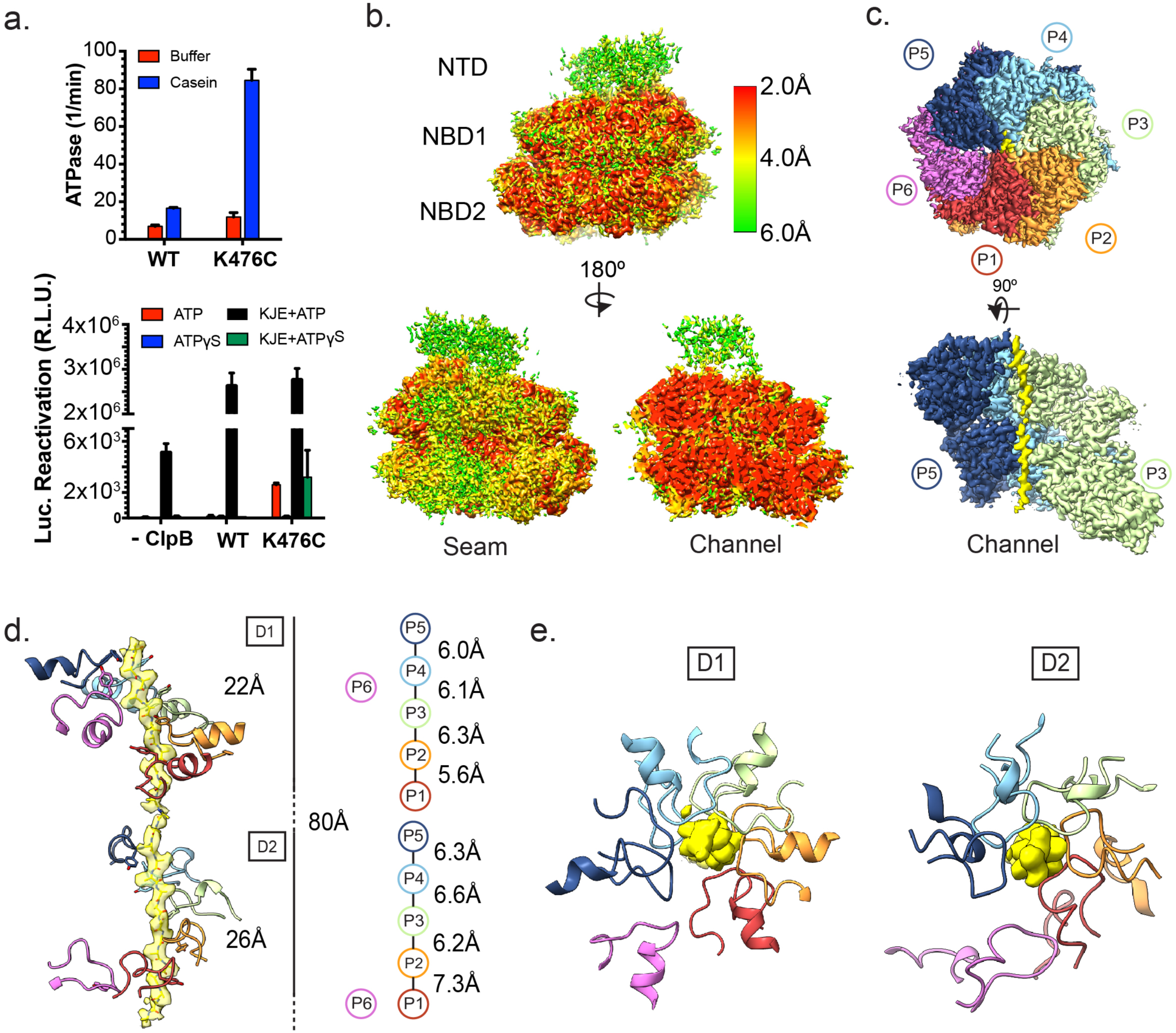
Activity and substrate-bound structure of the ClpB^K476C^ hyperactive variant. **(a)** ClpB and ClpB^K476C^ ATPase activity (upper panel) measured in the presence or absence of casein. Y-axis values are of phosphate release (in^−1^) and represent means±SD (n=2). ClpB and ClpB^K476C^ luciferase disaggregase activity (lower panel), as relative luminescence, measured in the absence or presence of KJE with either ATP or ATPγS. Values represent means ±SD (n=2).EM density map of the ATPγS-ClpB^K476C^:casein complex **(b)** colored by resolution^64^ and **(c)** colored to show individual protomers (P1-P6) and substrate (yellow). **(d)** Side and **(e)** top-views of the NBD1 and NBD2 Tyr-containing pore loops (colored by protomer) with substrate EM density (yellow) modeled with a 27-residue poly-Ala chain. A schematic is shown with the distances (Å) between Tyr-substrate contacts along the NBD1 and NBD2 for protomers P1-P5; protomer P6 (magenta) is disconnected from the substrate.

Compared to WT, ClpB^K476C^ displayed ~2-fold elevated ATPase activity in the absence of casein and ~5-fold elevated ATPase activity in the presence of casein (Figure 1a). Moreover, in the presence of ATP but without the Hsp70 chaperone system (DnaK, DnaJ, and GrpE [KJE]), ClpB^K476C^ displayed elevated luciferase disaggregase activity compared to WT (Figure 1a). Addition of KJE greatly stimulated luciferase disaggregase activity of ClpB^K476C^ and ClpB (Figure 1a). Indeed, under these conditions ClpB^K476C^ displayed similar activity to WT (Figure 1a). Replacing ATP with ATPγS in the presence of KJE eliminated ClpB disaggregase activity, whereas ClpB^K476C^ retained some disaggregase activity (Figure 1a). Previously, we established that neither ClpB nor ClpB^K476C^ can remodel seminal amyloid^41^. Nonetheless, these data establish that ClpB^K476C^ displays robust disaggregase activity in vitro, which is partially independent of KJE and less sensitive to inhibition by ATPγS.

Given the robust activity and stable substrate-bound complex formation with casein and ATPγS, ClpB^K476C^ was targeted for high-resolution structure determination by cryo-EM. The final map refined to an indicated 2.9 Å resolution using the total dataset following sorting by 2D classification (Supplementary Fig. 1c and Supplementary Table 1). Extensive 3D classification was performed (see below) but did not improve the resolution of the substrate channel. Local resolution estimation reveals that the channel and protomers bound to the substrate are at the highest resolution for the complex (<3.0 Å) while the seam protomers are at a lower resolution (~4.0 Å), indicating flexibility at this interface (Figure 1b). The angular distribution of the particles show some top and side-view preferred orientations, and the 2D projections of the map exhibit well-defined features that match the reference-free averages (Supplementary Fig. 1d,e). The protomers (P1-P6) adopt a right-handed spiral configuration similar to what has been previously described^5,6^, and well-defined polypeptide density is identified to span the 80 Å-length of the NBD1 and NBD2 that comprise the translocation channel (Figure 1c). This density is attributed to a 27-residue unfolded strand of the casein substrate (Figure 1d). The molecular model for the AAA+ domains was built and refined using PHENIX^42^, using model derived from the bacterial ClpB structure^7^ (Supplementary Fig. 1f) and the AAA+ subdomains show well-defined features indicative of the resolution (Supplementary Fig. 1g). The lower-resolution NTDs were classified and modeled separately (see below). To determine if the ClpB^K476C^ is in the same arrangement as WT, an additional data set was also collected of WT ClpB bound to casein and ATPγS (Supplementary Table 1) and refined to 4.1 Å resolution (Supplementary Fig. 1h). Comparison of the WT and ClpB^K476C^ maps revealed they are overall identical in conformation (Supplementary Fig. 1i, j)

In the ClpB^K476C^:casein structure five protomers directly contact the substrate, which is positioned slightly off center to the channel axis and closer to the back protomers (P2-P5), opposite the seam interface (Figure 1d, e). The conserved NBD1 and NBD2 Tyr-containing pore loops, which are required for translocation^3,16^, are each separated by ~6–7 Å along the substrate axis and rotate on ~60°, based on the position of the substrate-interacting Tyr (Figure 1d). This arrangement indicates an overall dipeptide spacing along the substrate similar to previous substrate-bound AAA+ structures^5,6^ and the conserved Tyr residues (Y251 for NBD1 and Y653 for NBD2) are identified to intercalate between substrate side chains and contact the backbone. The more flexible seam protomer, P6, which is adjacent to the highest (P5) and lowest (P1) contact positions is asymmetric compared to the P1-P5 spiral and disconnected from the substrate, with the NBD1 and NBD2 pore loops positioned approximately 4–5 Å away (Figure 1d, e). Notably, this 5-bound:1-unbound spiral arrangement of the protomers is similar to other substrate-bound AAA+ complexes, supporting a conserved substrate interaction mechanism^34–36^.

### Distinct substrate gripping interactions by the NBD1 and NBD2 pore loops

The resolution of the substrate channel is improved substantially compared to previous structures of ClpB and Hsp104, allowing for precise mapping of the pore loop interactions that stabilize the unfolded polypeptide substrate. Two distinct NBD1 loops (D1, residues 247–258 and D1’, residues 284–295) extend into the channel and contact the substrate (Figure 2a and Supplementary Fig. 2). In the canonical pore loop (D1), the conserved Tyr (Y251), which is the primary substrate-interacting residue^16^, is flanked by basic residues in ClpB and Hsp104 (K250 and R252 in ClpB and K256 and K258 in Hsp104) (Figure 2b). This pattern is distinct from the well-characterized aromatic-hydrophobic pair motif (Ar/ϕ) (e.g. Tyr-Val) found in other AAA+ translocases and domains, including NBD2 of ClpB and Hsp104^43^. In this structure, we identify that K250 and R252 extend perpendicular to the substrate axis and together with Y251, which intercalates between the side chains of the substrate, form a well-defined clamp around the polypeptide (Figure 2c). Remarkably, the side chains of K250 and R252 extend perpendicular and contact E254 and E256 in the neighboring lower and upper pore loops, respectively (Figure 2d). Based on the orientation and distance between the side chains, these interactions appear to be salt bridges that form across the pore loops in all protomers except at the spiral seam (with protomer P6) and likely stabilize the flexible pore loops in the presence of substrate and coordinate dipeptide spacing.

**Figure 2:**
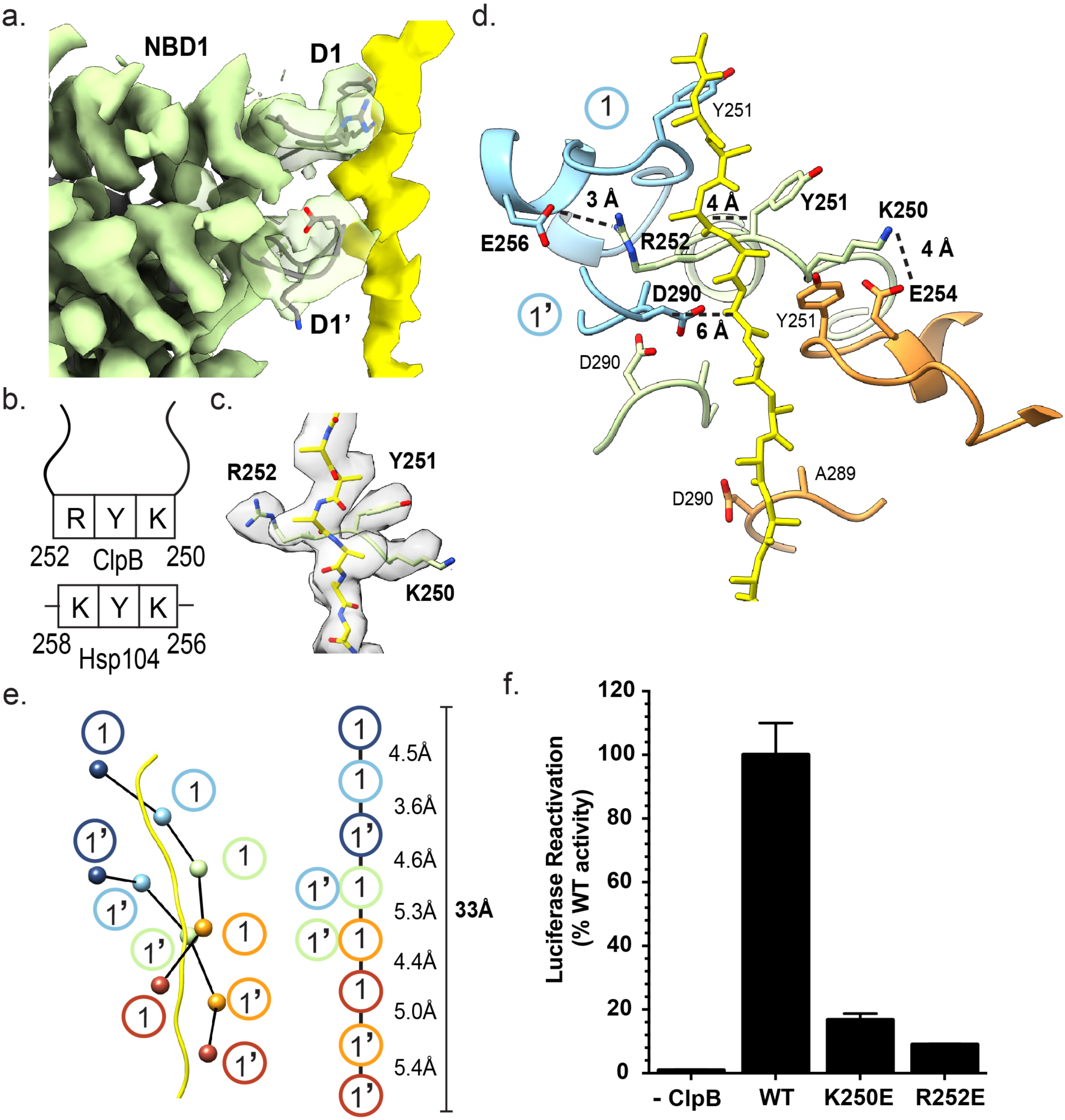
Substrate interactions by the NBD1. **(a)** Side view of the P3 NBD1 pore loop-substrate interactions with the D1 and D1’ loops indicated. **(b)** Schematic representation of the conserved D1 loop residues in ClpB and Hsp104. **(c)** Map and model of P3 D1 pore loop showing arrangement of K250, Y251 and R252 along the substrate density modeled with poly-Ala (yellow). **(d)** P2-P4 D1 and D1’ pore loops and modeled poly-Ala substrate with substrate interactions and proposed salt bridge interactions: E256-R252 and K250-E254 shown with approximate distances (dotted line). **(e)** Schematic of the double spiral of substrate interactions that is formed from the D1 and D1’ loops, with distances along the substrate axis shown based on the position of Y251 in the D1 and D290 in the D1’ pore loops (right). **(f)** ClpB, ClpB^K450E^, or ClpB^R252E^ luciferase disaggregase activity was measured in the presence of KJE with plus ATP. Values represent means±SD (n=2).

The secondary pore loops (D1’) are more flexible and variable around the hexamer, however A289 and D290 are consistently oriented toward the substrate and are approximately ~6 Å away from the backbone (Figure 2d). These loops form an additional spiral of substrate contacts that are shifted ~8 Å down the axis from the D1 loops (Figure 2e). Together the D1 and D1’ loops enable an ~11 amino acid-length (33 Å) of the polypeptide substrate to be stabilized by NBD1. While the D1’ loops increase the overall substrate interactions made by the NBD1, given the variability and distance from the substrate, the contributions are considerably smaller compared to binding by the KYR motif in the D1 loops.

The cross pore-loop interactions mediated by E256-R252 and K250-E254 salt-bridge interactions have not been observed previously and is surprising given the flexibility of the pore loops in the absence of substrate^7,8^. Therefore, the functional importance of these interactions was assessed by making K250E or R252E ClpB mutants. ClpB^K250E^ and ClpB^R252E^ were both defective in the ability to drive luciferase disaggregation and reactivation in the presence of KJE (Figure 2f). Indeed, ClpB^K250E^ exhibited ~17% WT activity, whereas ClpB^R252E^ exhibited ~9% WT activity (Figure 2f). These findings suggest that the integrity of the E256-R252 and K250-E254 salt bridges are important for ClpB disaggregase activity.

NBD2 similarly contains two pore-loop strands in each protomer that extend and contact the substrate polypeptide for P1-P5 (Figure 3a and Supplementary Fig. 3). These are the canonical loop (D2, residues 647–660) containing the conserved Ar/ϕ (Y653 and V654) motif and an additional short helical loop (D2’, residues 636–646). In the D2 loop, Y653 and V654 together form a clamp around the substrate backbone, intercalating between the side chains, appearing to make similar contributions to substrate binding (Figure 3b). Additional cross-pore loop interactions are not observed, thus the Ar/ϕ motif makes the primary substrate stabilizing for the NBD2. The Tyr residues are in a different orientation compared to the D1 residues, projecting more downward, towards the N-terminal face of the channel. This “up/down” configuration is similar to that identified for the Hsp104 structure^6^. For the D2’ loop residues E639, K640 and H641 project into the channel and appear to interact directly with the substrate polypeptide (Figure 3c). E639 and H641, in particular, are adjacent the side chains of the substrate at certain positions, with H641 positioned between the Y653 residues of two clockwise protomers. Thus, the D2’ loops form a well-defined additional spiral of stabilizing substrate interactions that coalesce towards the bottom of the NBD2 where the polypeptide exits from the translocation channel (Figure 3d).

**Figure 3:**
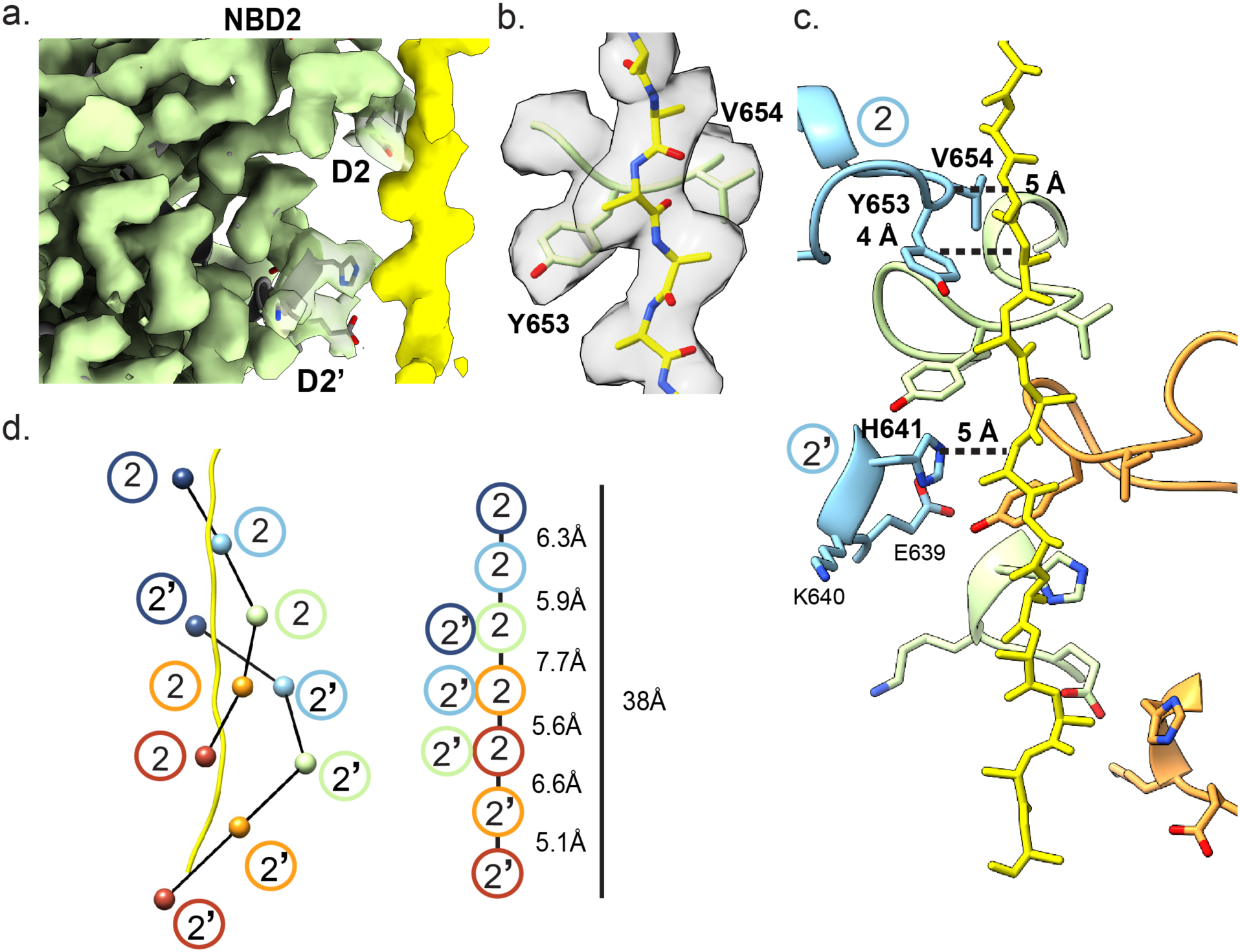
Substrate interactions by the NBD2. **(a)** Side view of the P3 NBD2 pore loop-substrate interactions with the D2 and D2’ loops indicated. **(b)** Map and model of P3 D2 pore loop showing arrangement of Y653 and V654, along the substrate density modeled with poly-Ala (yellow) **(c)** The D2 and D2’ pore loops of P2-P4, colored by protomer, and substrate (yellow), with distances between interacting residues and substrate backbone shown. **(d)** Schematic of D2 and D2’ loops interacting along the substrate for protomers P1-P5 with distances along the substrate axis shown based on the position of Y653 and H641 for the D2 and D2’, respectively.

Overall, the structure reveals distinct substrate-interaction mechanisms by the NBD1 and NBD2 pore loops. The D1 loops form strong substrate interactions involving direct contact by Y251 and flanking interactions by K250 and R252, which form additional salt bridge-stabilizing interactions with the adjacent pore loops. The substrate interactions by the D2 loop involve primarily a clamp-like configuration of the canonical Ar/ϕ motif around the polypeptide. The secondary, D1’ and D2’ loops also appear to play key roles in substrate binding; most notably for the D2’ E639 and H641 residues, which contact the substrate towards the channel exit.

### Substrate interaction specificity by the NBD1 and NBD2

The cryo-EM density corresponding to the casein substrate polypeptide is well-defined across the NBD1 and NBD2 portions of the channel (Figure 1c). This finding is surprising considering that ClpB can slowly hydrolyze ATPγS and translocate substrates under these conditions ^40,44^. The density for casein would thus be expected to heterogeneous and blurred out in the reconstruction. Instead, we hypothesize that ClpB may interact preferentially with certain sequences that may stall translocation, leading to the increased homogeneity and improved resolution of the polypeptide density we observe in the channel.

To model the polypeptide substrate in the structure, 1604 models of casein peptides were threaded into the density for the NBD1 and NBD2 rings using Rosetta^45^. The model of the lowest energy fits are shown (Figure 4a,b) and reveal favorable binding to casein sequences: VVTILALTLPF for the NBD1 and ILACLVALALA for the NBD2. Similar low energy scores were also achieved for additional peptides, while many peptides scored unfavorably as well, indicating preferential interaction with specific casein peptide sequences by the NBD1 and NBD2 (Supplementary Table 2). To characterize the specificity further, amino acid preferences at each position were ranked based on the threading analysis (Figure 4c,d). This was achieved by taking the optimal casein peptides (above) and individually mutating each of the residues to all 20 amino acids, then comparing the energetics of sidechain interactions at these mutated positions in arrangements consistent with the peptide backbone density. Energetically favorable interactions are identified primarily for large aromatic and hydrophobic residues for both the NBD1 and NBD2, while residues with small sidechains, such as Pro and Gly, interact unfavorably (Figure 4c,d). Particularly for NBD1, the lower energy, favorable interactions alternate along the peptide sequence in a manner corresponding approximately to the substrate residues that stack between the conserved Tyr residues in the pore loops. Notably, interactions by NBD1 are lower energy and more specific compared to the NBD2, likely reflecting the differences in the pore loop interactions discussed above (KYR for the NBD1 compared to YV for the NBD2) (Figure 4c,d). Overall these results reveal that ClpB readily accommodates bulky residues and interacts favorably with specific sequences containing aromatic and hydrophobic residues in an alternating arrangement that matches the pore-loop spacing.

**Figure 4.**
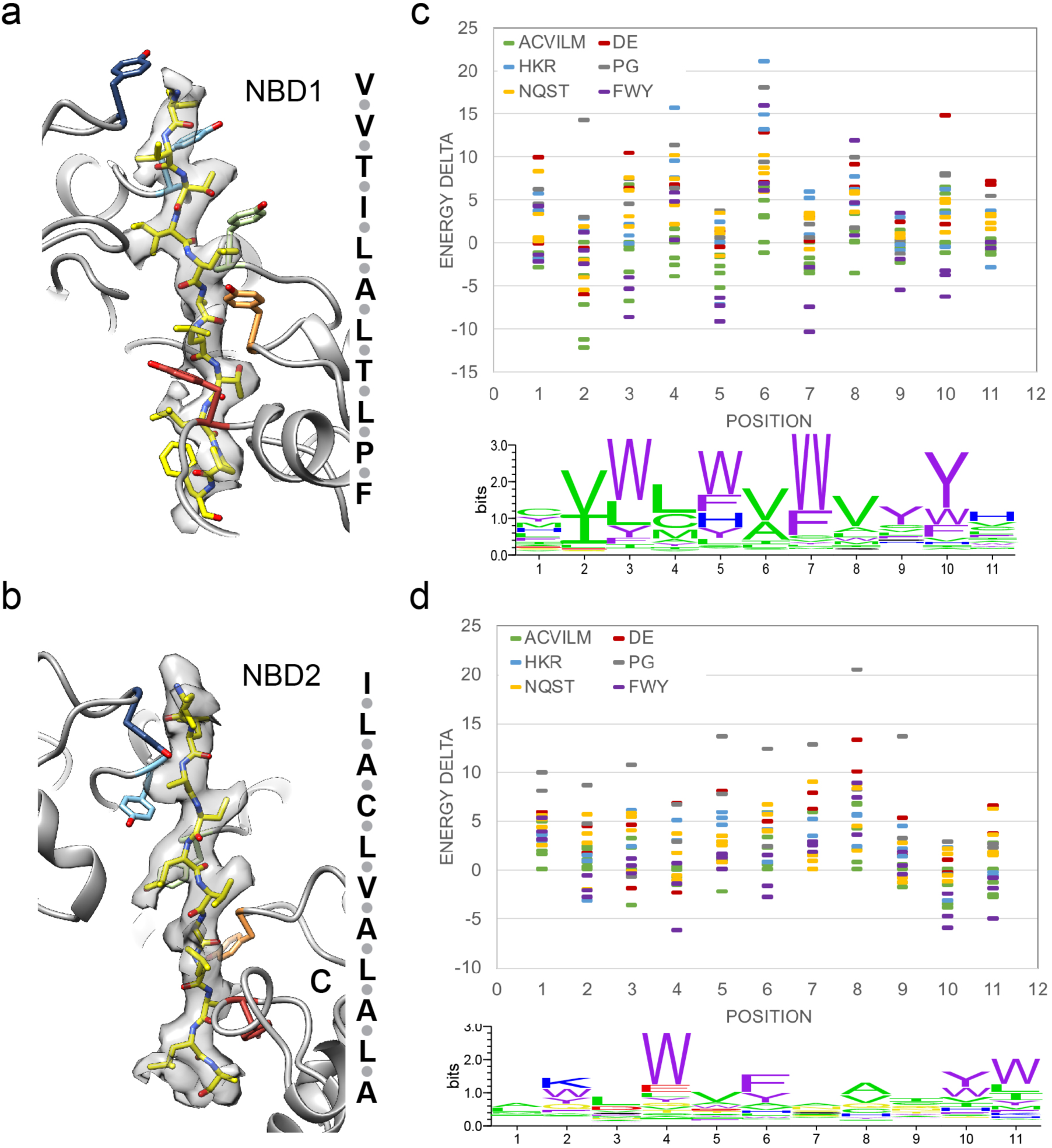
Analysis of the substrate sequence stabilized by the NBD1 and NBD2. Molecular model and sequence of the energetically favored casein peptide sequence (yellow) in density for the NBD1 **(a)** and NBD2 **(b)**. Interacting Tyr resides (Y251 and Y653) are colored by protomer. A plot showing per-energy deviations at each of the eleven positions interacting with NBD1 **(c)** and NBD2 **(d)**; the logo plots convert these energies to a distribution, showing the sequence preferences at each position.

### Focused refinement reveals NTD ring architecture and substrate engagement

The polypeptide substrate is expected to enter the AAA+ NBD1-NBD2 translocation channel via the NTDs^3^, which form an additional ring connected to the NBD1 ring^5,6^ (Figure 1b). However, resolution of the NTDs has been challenging due to their flexibility, thus substrate engagement at the channel entrance has been difficult to resolve. To better resolve the NTDs in the ClpB^K476C^:casein structure, particle re-centering and focused refinement was performed (Supplementary Fig. 4a). The density for the individual NTDs and connecting linker to the NBD1s were improved by this procedure (Supplementary Fig. 4b), enabling the identification of three NTDs that comprise the channel entrance (Figure 5a). The NTDs are identified to be associated with protomers P1, P3 and P5, revealing that interactions between alternating protomers together form an NTD ring and channel entrance which interacts with the polypeptide substrate and retains the overall spiral architecture of the NBDs (Figure 5a). Additional NTD density corresponding to the other protomers (P2, P4 and P6) is not observed, even at reduced threshold values, indicating they are likely disconnected and flexible (data not shown). Density corresponding to the polypeptide substrate is identified to extend up from the P5-NBD1 pore loop and directly contact the P5 and P3 NTDs (Figure 5b). An additional 9 residues were modeled into this substrate density as alanines, thereby bringing the total length of substrate to 34 residues that is stabilized in an unfolded arrangement across the three rings of ClpB. Based on the molecular model, NTD-substrate interactions involve helix A6, which contacts the substrate approximately 2 residues above P5-NBD1 pore loop interaction in protomer P3 and ~9 residues above in protomer P5 (Figure 5c). Interactions are also observed between helix A1 in protomer P3 and the substrate (Figure 5b,c). In support of these interactions, residues in helices A1 and A6 that appear to be adjacent the substrate polypeptide in the structure were previously identified to form a substrate binding-groove and interact with hydrophobic regions of substrates^32^ (Figure 5c). Overall, we identify the NTD ring is comprised of three domains from alternating protomers, P1, P3 and P5, which together form an additional right-handed spiral and directly contact the substrate. The identification of the extended polypeptide strand in the NTD ring contacting the P3 and P5 NTDs reveals that the casein substrate is stabilized in an unfolded state and suggests a direct role for the NTDs in substrate transfer to the AAA+ domains.

**Figure 5:**
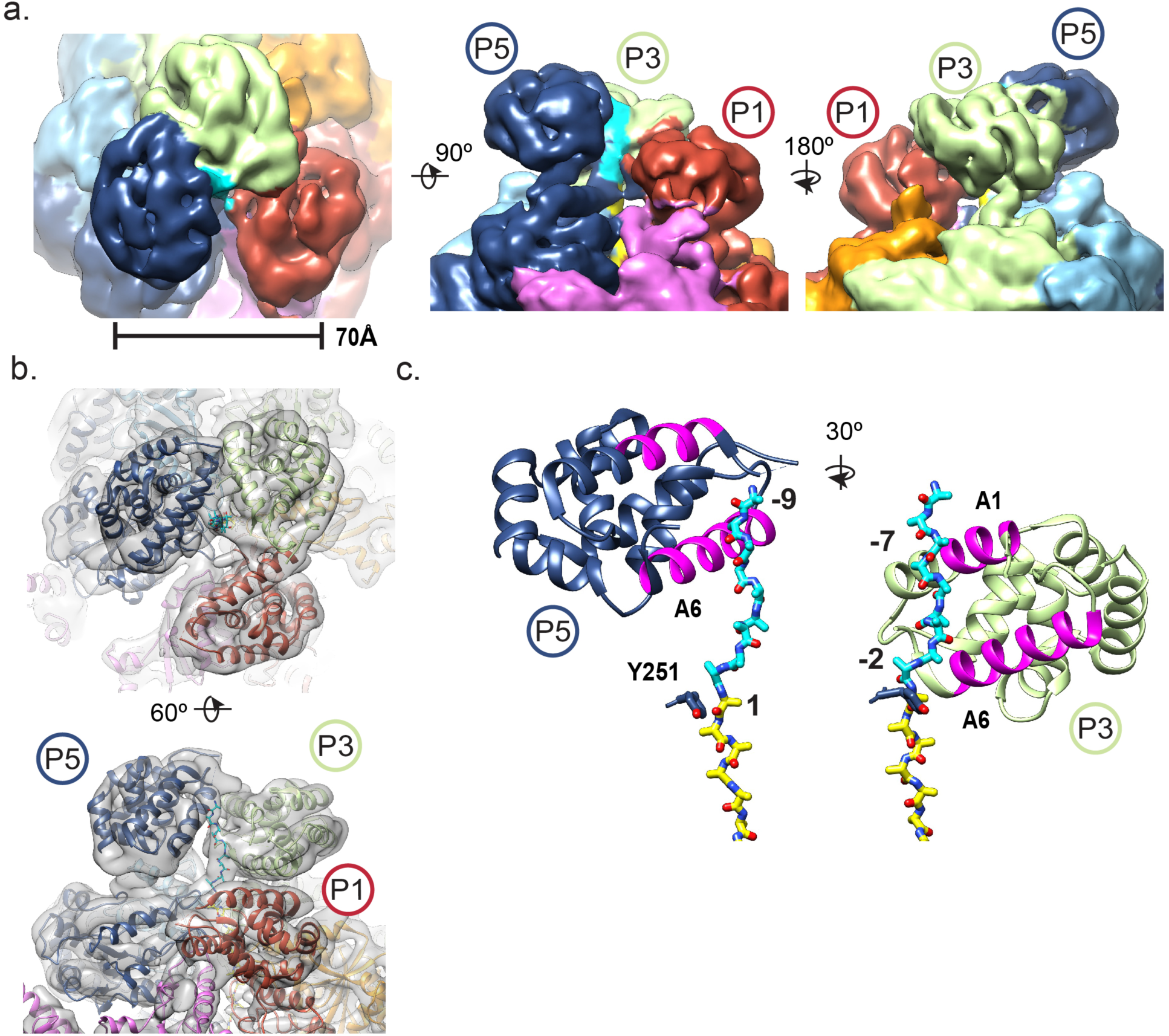
3D focus classification reveals substrate interaction with NTDs. **(a)** Views of the NTD-focused refinement map after particle re-centering, colored by protomer, showing the P1-P3-P5 trimeric NTD ring with the diameter indicated. **(b)** Map and molecular model of the P1-P3-P5 NTD trimer, colored by protomer, and substrate polypeptide, modeled as poly-Ala, that extends from the NBD1 (cyan). **(c)** Model of P5 and P3 NBDs interacting with substrate. Substrate interacting helices A1 and A6, previously characterized^32^, are shown.

### Distinct protomer conformations and nucleotide states at the spiral interface

The previous substrate bound structures identified that the protomers at the spiral seam are flexible and undergo conformational rearrangements. In particular, 3D classification revealed two conformations of Hsp104:casein that support a rotary mechanism involving substrate release and re-binding concurrent with ATP hydrolysis at this interface. Similarly, for the ClpB^K476C^ structure, the protomers at the spiral interface are identified to be flexible and at a lower overall resolution following refinement of the total dataset (after 2D classification) (Figure 1b). Following 3D classification two distinct conformations (Pre and Post states) of the seam protomers, P1 and P6, are identified and refined to 3.4 Å for the Pre-state and 3.7 Å for the Post state (Supplementary Fig. 5a). To improve the map of the P6-P1 interface, particles were re-centered and a 3D mask was applied during the refinement (Supplementary Fig. 5b and Figure 6a). The Pre-state P6-P1 protomer arrangement is identical to the model determined from refinement of the total dataset (discussed above) and similar to the Hsp104 closed state and ClpB^BAP^ structure (Supplementary Fig. 5c). For both states, protomer P6 remains disconnected from the substrate, unlike the “Extended” state identified for Hsp104 where all six protomers are bound in the spiral arrangement. Additionally, by comparison to the 20 models of the 3D classification (Supplementary Fig. 5a), the Extended conformation was confirmed to not be present in the dataset (data not shown), indicating ClpB^K476C^ does not adopt this conformation under these same casein-binding conditions.

**Figure 6:**
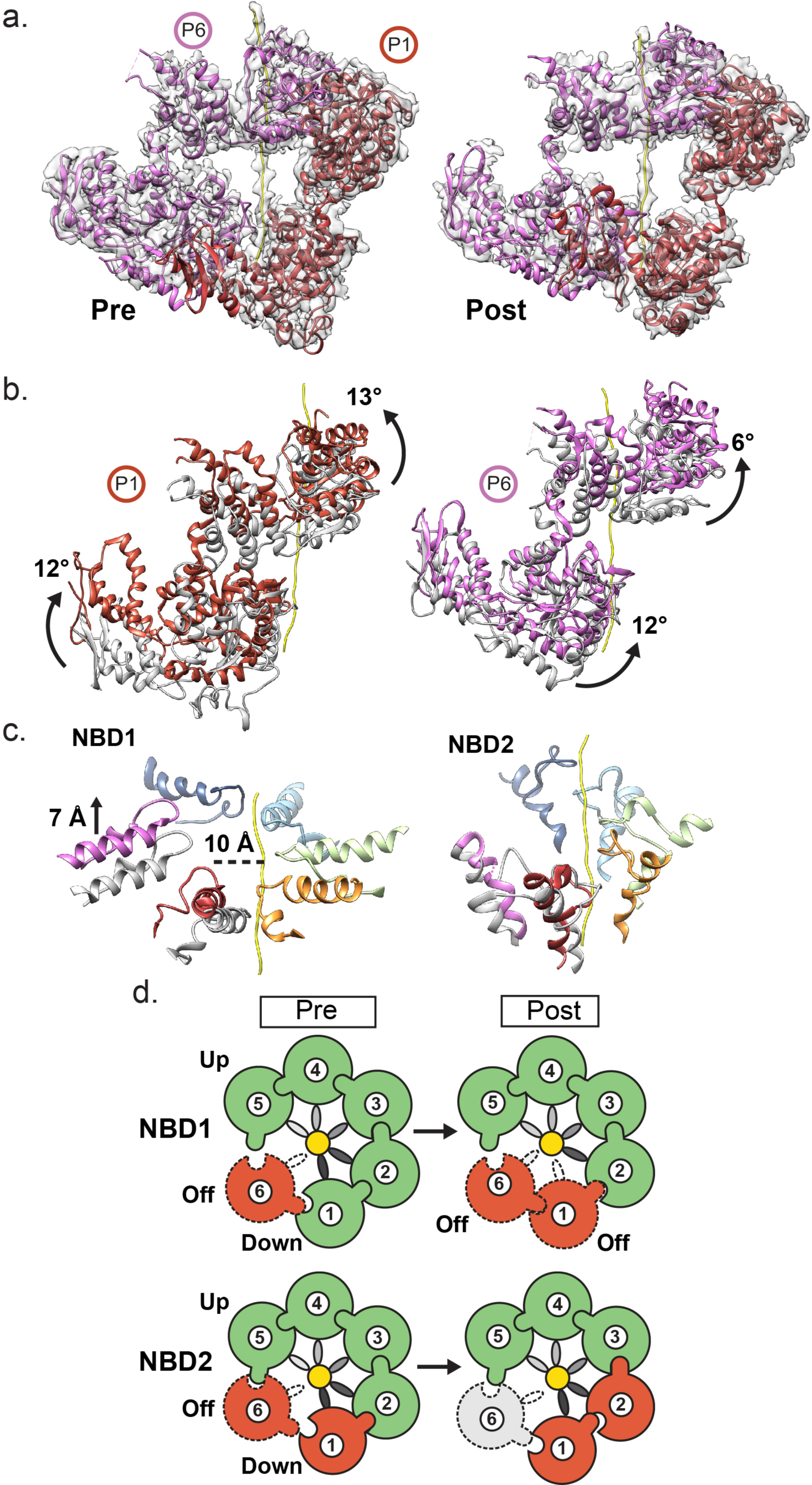
Conformational changes and nucleotide states of the protomers at the spiral seam. **(a)** Cryo-EM map and molecular model of protomers P1 and P6 in the “Pre” and “Post” conformational states following particle re-centering and focused refinement of the spiral seam. **(b)** Protomers P1 and P6 for both Pre (grey) and Post (colored) states with substrate (yellow), shown superimposed following alignment to protomer P3 in the hexamer. Conformational changes are shown as distinct rotations (arrows) of the NBD1 and NBD2. **(c)** The Pre- (grey) and Post-state (colored) NBD1 and NBD2 pore loops shown superimposed for protomers P1 and P6 with distances indicated following alignment of P3 pore loops in the hexamer for both states. **(d)** Schematic showing proposed NBD1 and NBD2 nucleotide states of the hexamer for the Pre and Post states, colored to indicate ATP (green), ADP (red), and APO (grey) state, based on analysis of the nucleotide pockets in the two structures (Supplementary Fig 6d). Protomers that are disconnected from the substrate are indicated with a dash and Arg-finger contact is indicated by the interlocking contact between two protomers.

The arrangement of the P6-P1 in the Post state, however, is distinct from the previous structures and involves conformational changes of the NBDs in both protomers (Figure 6b). In comparing Pre to Post state conformational changes, NBD1 of P1 and P6 rotate together clockwise and upwards along the substrate axis by approximately 13° and 6°, respectively (Figure 6b and Supplementary Movie 1). The NBD2s similarly rotate upward and towards the substrate axis in a clockwise manner. Most notably, the conformational changes alter the position of the NBD1 pore loops. The P1-NBD1 loop in the Post state shifts upward along the substrate axis by ~9 Å and rotates away, becoming disconnected from the substrate and asymmetric to the helical position of the P2-P5 pore loops (Figure 6c). The P6-NBD1 loop similarly shifts upward by ~7 Å, becoming nearly parallel with P5 at the top contact site, but remains disconnected from the substrate. Based on these comparisons, the Post state appears to be in an intermediate translocation state with respect to the NBD1 pore-loop positions, with protomer P6 on-path to bind next site along polypeptide, above the P5 position, while protomer P1 appears to be transitioning to a disconnected state, beginning with release by the NBD1 pore loop. Together, these conformational changes indicate that the P1-P6 protomers may simultaneously rearrange during the translocation step with P1 releasing substrate while P6 becomes repositioned to bind substrate at the next contact site above P5.

The nucleotide pockets were compared to further characterize these states. The NBD1 pockets for protomers P2-P5 in both Pre and Post states show well-defined density for ATPγS and the Arg finger (R331)-γ-phosphate interaction is identified, indicating an active, ATP state configuration (Supplementary Fig. 5c). Additionally, the P1-NBD1 is also confirmed to be in an ATP-active state for the Pre-state conformation, based on the nucleotide density and position of the Arg finger (Supplementary Fig. 5d). However, in the Post state, the P1-NBD1 appears to be in a post-hydrolysis, ADP state and inactive based on the P5 Arg finger position. Additionally, the P6-NBD1 for both the Pre and Post states shows reduced density for nucleotide and appears to be in a post-hydrolysis state (Supplementary Fig. 5d). Although, density in this region is not well-resolved for the Post state. For the NBD2, protomers P3-P5 are in an ATP-active state and bound to ATPγS which is contacted by the Arg finger (R756) from the clockwise protomer by for both the Pre and Post conformations (Supplementary Fig. 5c). For the P1 and P6 protomers the nucleotide pockets appear to be largely in an ADP state for both the Pre and Post conformations, however the P6-NBD2 is likely in apo state, based on the overall reduced density for a bound nucleotide (Supplementary Fig. 5d). Notably, the P2-NBD2, which is bound to the next position along the substrate compared to P1, appears to switch from an ATP-active state to an ADP state between the Pre and Post conformations, indicating hydrolysis may occur at this site during the conformational change (Supplementary Fig. 5d). In summary, the stable, substrate-bound protomers (P2-P5) remain largely in an ATP-active state, while the protomers at the spiral interface are predominately in an ADP state (Figure 6d). Sites at P1-NBD1 and P2-NBD2 shift from ATP to ADP states during the Pre to Post conformational change, supporting a mechanism whereby hydrolysis occurs as protomers transition to the lower contact sites (P1) and substrate-release state (P6) at the seam interface (Figure 6d).

### Hexamer asymmetry reveals expansion between the NBD1 and NBD2 rings coincides with substrate release

In side views of the reference-free 2D projection averages the NBD1 and NBD2 rings appear to adopt a non-parallel configuration identified by an increase in the distance between the rings on one side versus the other (Figure 7a). This asymmetry is further identified in both Pre and Post-state hexamer models, in which the overall width of the NBD1-NBD2 double ring increases at the P6-P1 spiral seam compared to the P3 and P4 protomers across the hexamer, going from ~75 to 106 Å in the Pre-state and ~77 to 96 Å in the Post-state (Figure 7b). Thus, in addition to the right-handed spiral arrangement of the pore loop-substrate interactions, which involve a ~6–7 Å shift per substrate contact site, the distance between the NBD1 and NBD2 rings increases moving down the substrate axis from P5 to the P6-P1 interface. Both the Pre and Post states show similar changes around hexamer. When the protomers are separated and individually aligned to the NBD1 large domain this expansion between the NBD1 and NBD2 can be readily observed and involves rotations about the NBD1-NBD2 connecting residues (545–555) (Figure 7c). In particular, the distance between residues I546 and P594 at the NBD1-NBD2 interface increases from 13 Å to ~30 Å going from the P5 protomer clockwise around the hexamer to P1 and P6. Considering that the pore loop contacts for both the NBD1 and NBD2 remain in a defined register along the substrate, this increased separation between the NBD1 and NBD2 may add strain on the pore loop-substrate interactions moving down the channel which could facilitate ATP hydrolysis and substrate release at the lower interaction sites.

**Figure 7:**
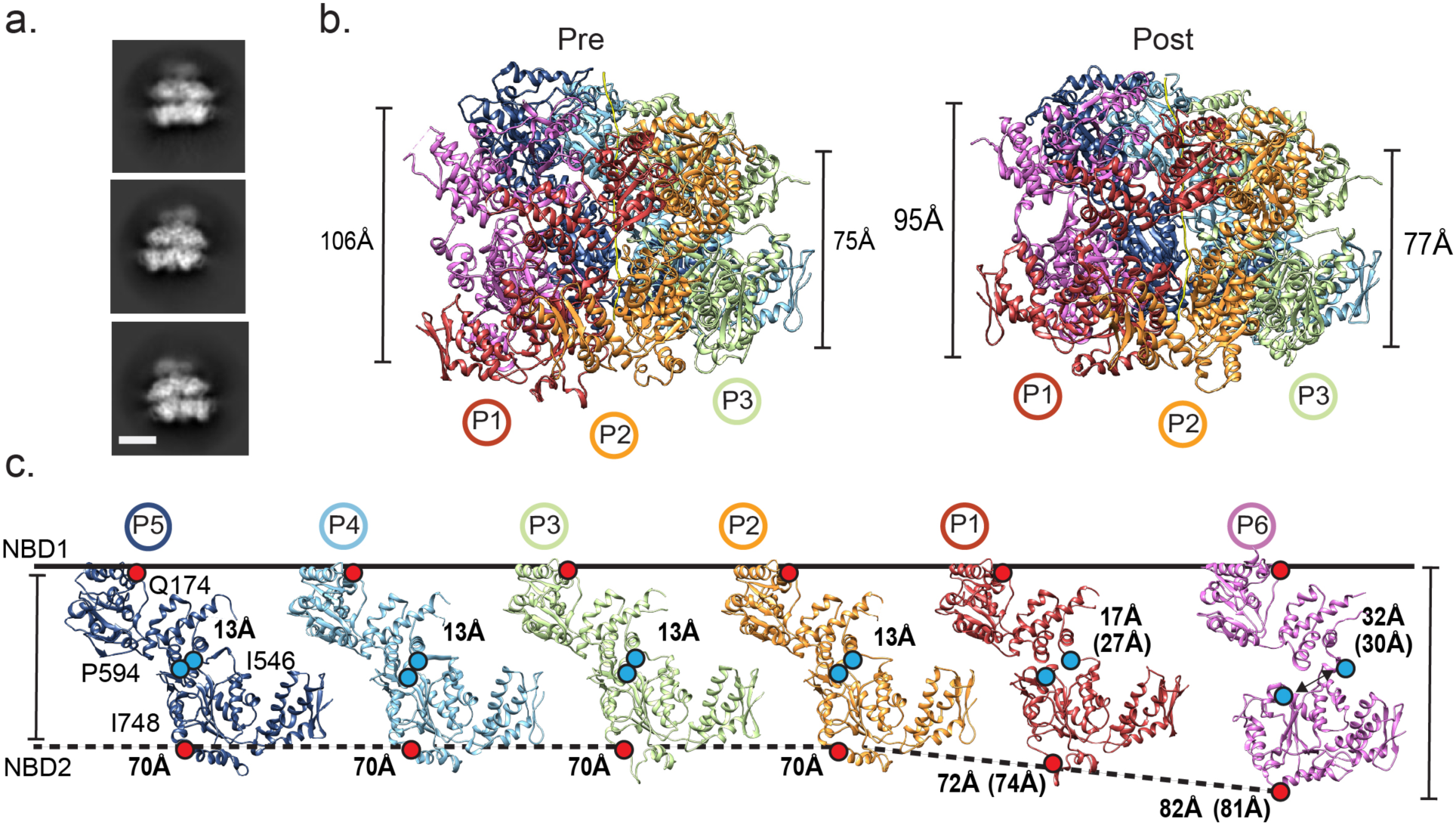
Expansion between NBD1 and NBD2 towards the seam interface. **(a)** 2D reference-free class averages of hexamer side views showing nonparallel arrangement of the NBD1 and NBD2 AAA+ rings. The scale bar equals 50 Å. **(b)** Side view of the Pre and Post-state models showing the expansion of NBD1-NBD2 rings measured at the P1-P6 spiral seam (left) compared to the P3-P4 interface across the hexamer (right). **(c)** The individual protomers from the Pre-state shown separated and aligned to the NBD1. Distance measurements are shown for residues Q174 and I748 (red dots) to depict overall protomer changes, and for residues I546 and P594 (blue dots) to show changes at the NBD1-NBD2 interface. Distances for the P1 and P6 protomers in the Post state are shown in parentheses.

## DISCUSSION

A number of recent structures of substrate-bound AAA+ translocases have begun to reveal a conserved mechanism of substrate interaction and translocation. For the Hsp100 disaggregases, key questions have remained about the specific roles of the two AAA+ domains, NBD1 and NBD2, and how conformational changes and ATP hydrolysis are coordinated around the hexamer to drive polypeptide translocation. To better elucidate the Hsp100 disaggregation mechanism we determined cryo-EM structures of the ClpB hyperactive variant, K476C, bound to the model substrate, casein. We focused on this variant, in particular because of its robust ATPase and disaggregase activity in order to capture active states of translocation (Figure 1a). Additionally, we found that the cryo-EM structure of this variant bound to casein went to reasonably high-resolution (2.9 Å resolution for the total dataset, Supplementary Fig. 1c and Table 1) whereas the WT complex appeared more heterogeneous and refined to a lower resolution but with an identical overall structure (Supplementary Fig. 1h,i). In the ATPγS-ClpB^K476C^:casein structure we identify a well-resolved translocation channel with a defined NBD1-NBD2 spiral of pore-loop-substrate contacts, revealing distinct substrate gripping mechanisms and sequence interaction specificity for the for the AAA+ domains. We identify a substrate channel entrance that is comprised of an NTD trimer that binds substrate as a spiral, extending its unfolded state, and thereby facilitating transfer to the AAA+ channel. Two conformations of the seam protomers are identified in different nucleotide states, revealing a translocation-step mechanism. Analysis of the hexamer arrangement additionally reveals an expansion between NTDs at the spiral interface that may contribute to substrate pulling and ATP hydrolysis-driven release during translocation steps.

The NBDs of ClpB, Hsp104 and related Hsp100s are members of distinct clades of the AAA+ family. The NBD1 is a member of clade 3 which includes FtsH, p97, NSF, and katanin, while NBD2 is a member of clade 5 which includes ClpX, RuvB and Lon^43,46^. Although evolutionarily distinct, the pore loops across these clades primarily contain the conserved Ar/ϕ substrate interacting motif, which is often flanked by Gly residues (e.g. Gly-Tyr-Val-Gly in *E. coli* ClpX, and the NBD2 of ClpB), indicating conserved functions^14^. The KYR motif, in which the substrate-interacting Tyr is flanked by basic residues, appears to be a unique feature of the NBD1 in disaggregases such as ClpB, Hsp104 and ClpA. Here we identify that K250 and R252 serve key roles in stabilizing the pore loops via salt bridge interactions with E254 and E256 in the pore loops of adjacent protomers (Figure 2d). Importantly, we identify that charge reversal mutations of either K250 or R252 results in a loss of activity *in vitro*, indicating these interactions are required for disagregase function. Together with Y251, which we identify to intercalate between the sidechains of the substrate (Figure 2c), these interactions likely contribute greater substrate binding energy compared to the Ar/ϕ motif of NBD2. Indeed, in the substrate modeling experiments we identify a strong substrate specificity for the NBD2 compared to the NBD1. Notably, Y251A has a less sever phenotype than Y653A^3^. Based on the structure here, the K250 and R252 cross pore-loop interactions may partially compensate for this loss of function in the NBD1 compared to the NBD2. Considering the NBD1 makes the first contact with substrate beyond the more flexible NTD, these strong interactions may be critical for initiating substrate unfolding or facilitating processivity.

In the NBD2, interactions across the pore loops are not observed, however Y653 and V654 form a well-defined clamp arrangement around the substrate backbone (Figure 3b). Based on our modeling experiments, the substrate sequence preferences for the NBD2 also favor aromatic and hydrophobic residues for positions that match the pore loop interactions. However, the energy change is not as substantial (Figure 4c,d), indicating the NBD2 exhibits weaker, and more nonspecific interactions compared to the NBD1. Thus, the NBD1 and NBD2 have distinct mechanisms for substrate gripping that likely reflect specific roles in disaggregation. Notably, we also identify contacts by secondary pore loops (D1’ and D2’), supporting roles for these additional residues in the channel in further stabilizing the substrate in an unfolded arrangement. In particular, we identify residues H641 and E639 in the lower portion of the NBD2 channel comprise the majority of contacts where the substrate would exit translocation channel. These weaker contacts may be important for substrate release.

The specificity for substrate sequences that contain hydrophobic residues agree with previous studies identifying ClpB binds preferentially to peptides containing aromatic and basic residues^16^. Here we identify preferences for aromatic and hydrophobic substrate residues, which are mediated by the canonical pore-loop residues and are particularly strong in NBD1, likely due to the KYR motif. Additionally, our results indicate that sequences containing stretches of residues with small sidechains, such as Gly, Ala and Pro, would be more unfavorable and potentially inhibitory to translocation by ClpB, or involve alternate substrate interaction mechanisms. Indeed, diversity in the substrate sequence is favored by the proteasome^47^ and Gly-Ala repeat sequences impair unfolding and trigger release, a mechanism by which the Epstein Barr virus protein EBNA1 evades proteasomal degradation^48,49^.

Our analysis of the more flexible NTD ring surprisingly reveals that the substrate entrance channel is comprised of an NTD trimer from alternating protomers (P1-P3-P5). Density for the remaining NTDs was not observed despite extensive classification, indicating these domains are likely flexible. In the WT Hsp104:casein structure density for all six NTDs was identified and localized to the NTD ring^6^, while in the ClpB^BAP^:casein structure a trimeric NTD arrangement was identified for one of the classes, supporting the arrangement observed here^5^. While the alternating three-protomer configuration is surprising, two isoforms, a full-length (ClpB95) and an NTD-minus ClpB (ClpB80), are present in *E. coli* and have been shown to form heteroligomers and function synergistically in disaggregation^50^. Therefore, ClpB may have evolved to function optimally as a heterohexamer with an NTD ring that consists of primarily three protomers which dynamically interchange to bind substrate during cycles of NBD-driven ATP hydrolysis and translocation. By contrast, Hsp104^ΔN^ subunits reduce prion dissolution by Hsp104 hexamers^33^. Importantly, our refinements of the ClpB NTD ring enabled modeling of the NTDs and substrate that identify the polypeptide extends from the AAA+ channel at the P5-pore loop position and contacts the known substrate-binding hydrophobic groove^32^ in protomers P3 and P5 (Figure 5c), revealing that the NTDs contribute directly to substrate translocation.

Based on our analysis of the Pre and Post states, we propose polypeptide translocation by ClpB involves a counterclockwise rotary mechanism whereby ATP hydrolysis drives substrate release at the lower positions along the substrate (P1 and P2 sites) and subsequent ATP binding enables substrate re-binding to the next contact site, above protomer P5, resulting a processive translocation step (Figure 6). This counterclockwise rotary cycle is similar to models proposed for Hsp104^6^ and other substrate-bound AAA+ structures^34,35^. By comparison of the Pre and Post conformational states, we identify that the NBD1 of the off protomer shifts upward along the substrate axis and on-path to the next contact site (Figure 6c), while the P1-NBD1 becomes unbound from the substrate, and thus on-path to the substrate release state. Additionally, the P2-NBD2 also appears to undergo hydrolysis during this conformational change, based on the ADP and ATP states of the nucleotide pockets of the Pre and Post states, respectively. ATP hydrolysis by NBD1 and NBD2 is cooperative and allosterically controlled^51,52^. The conformational changes we identify here indeed support ATP hydrolysis-driven coupling between the NBD1 and NBD2 and indicate this regulation may function to destabilize substrate interactions at the lower substrate contact sites to facilitate translocation.

Finally, we identify an increase in the distance between the NBD1 and NBD2 in the protomers positioned lower along the substrate and at the spiral seam compared to the upper, substrate bound protomers (Figure 7c). Considering the pore-loop contacts remain evenly spaced along the substrate, this NBD1-NBD2 separation could function as a mechanical “pulling” mechanism that maintains the unfolded polypeptide in the channel and coordinates hydrolysis to occur at the lower substrate-interaction sites where the distance between the NBD1 and NBD2 is greater. Additionally, we find that the P6 protomer, which is unbound to substrate and in a post-hydrolysis ADP or apo state for both NBDs, has the largest separation between the NBD1 and NBD2, and therefore is likely in a ground-state or relaxed configuration. Based on these conformational differences we propose the NBD1 and NBD2 function together as a spring that becomes set upon ATP and substrate binding in a compact state. As protomers shift to lowest position along the substrate, the separation between the NBD1 and NBD2 increases, adding strain to these interactions, thereby triggering ATP hydrolysis and substrate release. In this manner, mechanical coupling of the NBD1 and NBD2 could be critical for processive translocation by directing stepwise substrate release and re-binding events to occur sequentially at the spiral seam.

## Methods

ClpB^K476C^ (150 μM) was incubated with ATPγS (5 mM) in the presence of FITC-casein (150 um) (#C0528, Sigma) before completing size exclusion chromatography (SEC) analysis and complex purification. The buffer used in the complex formation contained: 40 mM Hepes (pH=7.5), 40 mM KCl, 10 mM MgCl_2_, 1 mM DTT. Samples was applied to a Superose 6 Increase 3.2/300 column (GE Healthcare) using the same buffer used in the incubation. Fractions were collected and analyzed by SDS-PAGE to ensure both ClpB^K476C^ and casein were present. The complex peak eluted at 1.5 mL (Supplementary Figure 1a). To ensure the complex remained stable during EM grid preparation, an additional 1 mM of nucleotide was added. The same procedure above was completed for the ClpB-ATPγS-casein complex.

### Activity analysis of ClpB

ClpB, ClpB^K476C^, ClpB^K250E^, and ClpB^R252E^ were purified as described^26,41^. For ATPase assays, ClpB (0.25μM monomer) WT or mutant was equilibrated in luciferase refolding buffer (LRB, 25 mM HEPES-KOH, pH 7.4, 150 mM KAOc, 10 mM MgAOc, 10 mM DTT) for 15 min on ice and then incubated for 5 min at 25°C in the presence of ATP (1 mM) plus or minus casein (10μM). ATPase activity was assessed by the release of inorganic phosphate, which was determined using a malachite green phosphate detection kit (Innova). Background hydrolysis was determined at time zero and subtracted. For luciferase disaggregation assays, aggregated luciferase was first generated by incubating firefly luciferase (50 μM) in LRB plus 8 M urea at 30°C for 30 min. The sample was then rapidly diluted 100-fold into LRB. Aliquots were snap frozen and stored at −80°C until use. Under these conditions, luciferase forms aggregated structures that are predominantly between ~500–2,000 kDa in size. These aggregates are detergent soluble and completely resolved by 1% SDS at 25°C. Aggregated luciferase (100 nM) was incubated with ClpB or ClpB^K476C^ (1 μM monomer), DnaK (1 μM), DnaJ (0.2 μM), and GrpE (0.1 μM), plus ATP regeneration system (5 mM ATP, 1 mM creatine phosphate, 0.25 μM creatine kinase) or 5 mM ATPγS for 90 min at 25°C. At the end of the reaction, luciferase activity was assessed with a luciferase assay system (Promega). Recovered luminescence was monitored using a Tecan Infinite M1000 plate reader.

### Cryo-EM Data Collection and 3D Reconstruction

The fractions collected after SEC were diluted to ~1.5 mg/mL and applied to glow-discharged holey carbon grid (R 1.2/1.3, Quantifoil). A vitrobot (Thermo Fischer Scientific) was used to plunge freeze the grids. Samples imaged on a Titan Krios TEM operated at 300 keV (Thermo Fisher Scientific). Images were recorded on a K2 Summit detector (Gatan Inc.) operated in super-resolution mode at 48,450X, which corresponded to a calibrated pixel size of 1.032 Å/pixel. Data was collected automatically using Serial-EM in which eight second exposures at 200 msec/frame, with a total electron dose of 56 e-per micrograph in 40 frames. MotionCor2 was used to perform whole-frame drift correction. During this process the first two frames of data were removed due to large amounts of drift and the frames were Fourier-binned by 2. Micrographs were CTF corrected using Gctf^53^ and poor micrographs were removed based upon visual inspection and CTF values. Single particles were picked automatically using Gautomatch with the standard picking conditions which amounted to 778,521 particles from 8,499 micrographs. Data processing was continued in cryoSPARC^54^. 2D classification was performed to remove contamination and junk particles; this amounted to about 8% of the dataset. An ab-initio model was created from the original particle set, which was then used in subsequent 3D classification and 3D refinement steps. The initial homogenous refinement of all 700K particles yielded a 2.9 Å resolution structure based on the .143 Fourier shell correlation criterion. From this structure the front two protomers, P1 and P6, and the NTDs were less resolved compared to the rest of the structure. To explore any changes that might be occurring to these areas, extensive 3D classification and focus classification was completed.

In order to perform the extensive 3D classification and focus classification steps, the data was processed using Relion 2.1^55^. An extensive 20-class classification was performed to determine if two different orientations of the mobile protomers could be uncovered. The original ClpB^K476C^ refinement map was used as the initial model after being low passed filtered to 60 Å. From this 3D classification, two different states were discovered, the Pre and Post states. The Pre-state (239,000 particles) was in the same orientation as the original 3D refinement map. The Post state (213,000 particles) had a different orientation around the mobile protomers, P1 and P6, with the stable protomers, P3-P5, staying the same. The different classes belonging to each state were grouped together and 3D refinement was performed. Approximately 248,000 particles were classified into junk particles classes and they were not further analyzed. Following 3D refinement the pre-state was refined to 3.4 Å and the post state was refined to 3.7 Å.

Two separate focus classifications were performed to improve resolution of the mobile protomers (P1 and P6) and the NTDs. A previous refinement run data file was used to re-center of the particles to the two mobile protomers or in the center of the NTDs (Supplementary Figure 5a and 6b). The particles were then re-extracted and re-centered and 3D classification was performed without image alignment with the reference map and a reference mask of the two front protomers or NTDs, both adjusted with the new center. For the mobile protomer focus classification, different classes were classified out matching the previous Pre and Post maps identified. The classes that provided improvement in resolution to the focus area were selected for 3D refinement, which included ~300,000 particles for pre-state and ~91,000 particles for post-state. The classes belonging to each state were grouped together and refined separately. For NTD focus classification, the classes that provided improvement in resolution to the focus area were selected for 3D refinement, which included ~272,000 particles. During 3D refinement the initial angular sampling value was set to the same value as the local search value. The final structure after focus classification provided an improvement in both the density and resolution of the front two protomers and the NTDs compared to the complete refined dataset map. All of the final maps underwent “Post-processing” procedures within RELION^56^ or cyroSPARC^54^ to sharpen and determine FSC estimation. ResMap^57^ was used to estimate the local resolution.

For the WT ClpB dataset, the fractions collected after SEC were diluted to ~1.5 mg/mL and applied to glow-discharged C-flat holey carbon grids (CF-2/1-4C-T, Protochips). A vitrobot (Thermo Fischer Scientific) was used to plunge freeze the grids and then they were imaged on a Titan Krios TEM operated at 300 keV (Thermo Fisher Scientific). Images were recorded on a K2 Summit detector (Gatan Inc.) operated in counted mode at 50,000X, which corresponded to a calibrated pixel size of 1.0 Å/pixel. Data was collected automatically using Leginon in which eight second exposures at 200 msec/frame, with a total electron dose of 52 e-per micrograph in 40 frames. MotionCor^58^ was used to perform whole-frame drift correction. Micrographs were CTF corrected in Relion using CTFFIND4^59^. Appion template picker^60^ was used to automatically pick single particles using templates from previously published Hsp104:AMPPNP dataset^20^. The total number of single particles picked before cleaning up in 2D classification was approximately 310,000. The rest of the data processing occurred in cyroSPARC^54^. 2D classification was used to remove junk particles and approximately 110,000 particles were then used in 3D classification and refinement. Using ab-initio reconstruction, an initial model was created which was then used in heterogenous refinement into 3 classes using the total particle set after 2D. The best refined class which contained 55,000 particles underwent homogenous refinement which refined to a resolution of 4.09 Å.

### Molecular Modeling

The initial model began by rigid body fitting previous hexamer ClpB model (pdb:5ofo)^5^ using UCSF Chimera’s^61^ *Fit in Map* function. The MD residues (409–524) were truncated from the model due to them not resolving in our structure. Missing D1’ loop residues were built de novo using Coot^62^. The substrate density within the channel was built de novo using Coot^62^ as a chain of alanines. Using real-space refinement in Phenix, the docked model underwent one round of simulated_annealing followed by 10 cycles of real-space refinement that consisted of minimization_global, rigid_body, adp, secondary structure restraints, and NCS. The resulting model then under-went manual real-space refinement in Coot^62^. To adjust and fix model issues identified through Molprobity validation, the model underwent another 10 cycles of real-space refinement in Phenix with only minimization_global, rigid body, and adp. For the Post-state model, 5ofo was docked in using UCSF Chimera’s^61^ Fit in Map function. The NBDs of P1 and P6 were then manually adjusted. The MD residues were truncated same as the pre-state model. The D1’ loops were not manually built due to missing density. The same procedure as the Pre-state was completed in Phenix.

For the casein sequence modeling, a backbone model was initially built manually corresponding to the density. Using Rosetta^45^, a “symmetrized” model was then generated by sampling four parameters corresponding to the phi/psi backbone angles of residues 1 and 2 of the two-residue repeat unit, and extending it over the 13 resolved residues in both the NBD1 and NBD2 rings. The phi and psi angles were sampled at 2-degree increments; each was evaluated against the density in both NBD1 and NBD2 rings, and a final model with phi_1_=−92, psi_1_=118, phi_2_=−102, and psi_2_= 110 was selected. Symmetrizing the model this manner was used to avoid overfitting the backbone model to the modest-resolution density.

Next, using the *partial_thread* tool in Rosetta, 1604 models of the casein sequence were threaded onto this backbone, 802 into the NBD1 and 802 into the NBD2 ring. Each of these models was then refined using Rosetta’s *relax* protocol with an additional term enforcing agreement to density at a modest weight (ensuring the backbone of the peptide would not refine far from the starting point). Rosetta’s *relax* allows refinement of both backbone and sidechain residues^63^; in this case, relax was restricted to only allow movement of the casein peptides themselves and the pore loops of NBD1 and NBD2, for a total of about 112 moving residues in each refinement. NBD1-and NBD2-interacting peptides were modeled separately.

For each of the 1604 threaded models, 5 independent refinement trajectories were carried out, and the lowest-energy model was selected for analysis. From these sequence-energy pairs, a profile was constructed by computing – for each amino-acid at each position – the average energy over all sequences with the corresponding amino acid at the corresponding position. The N- and C-terminal residues were excluded from this analysis. To make energy comparisons valid between diverse sequences, a sequence-specific reference weight was used in Rosetta to capture unfolded state energetics.

Finally, a “null-background” model was constructed, in which every residue except one was modelled as alanine: at this residue, all 20 amino-acid identities were generated. This led to a total of 440 models (11 positions x 20 amino-acid identities x 2 rings), which were refined in the same manner as the casein peptides. For these models, a profile was constructed by comparing the energy of each position-residue pair to the average energy of all evaluated peptides. These profiles showed similar trends to the casein-threaded profiles (data not shown).

## Data Availability

For the Pre- and Post-state ClpB reconstructions cryo-EM maps and atomic coordinates will be deposited to the Protein Data Bank. All other data are available upon request to the corresponding author.

## Supplementary Figures

**Supplementary Fig. 1:**
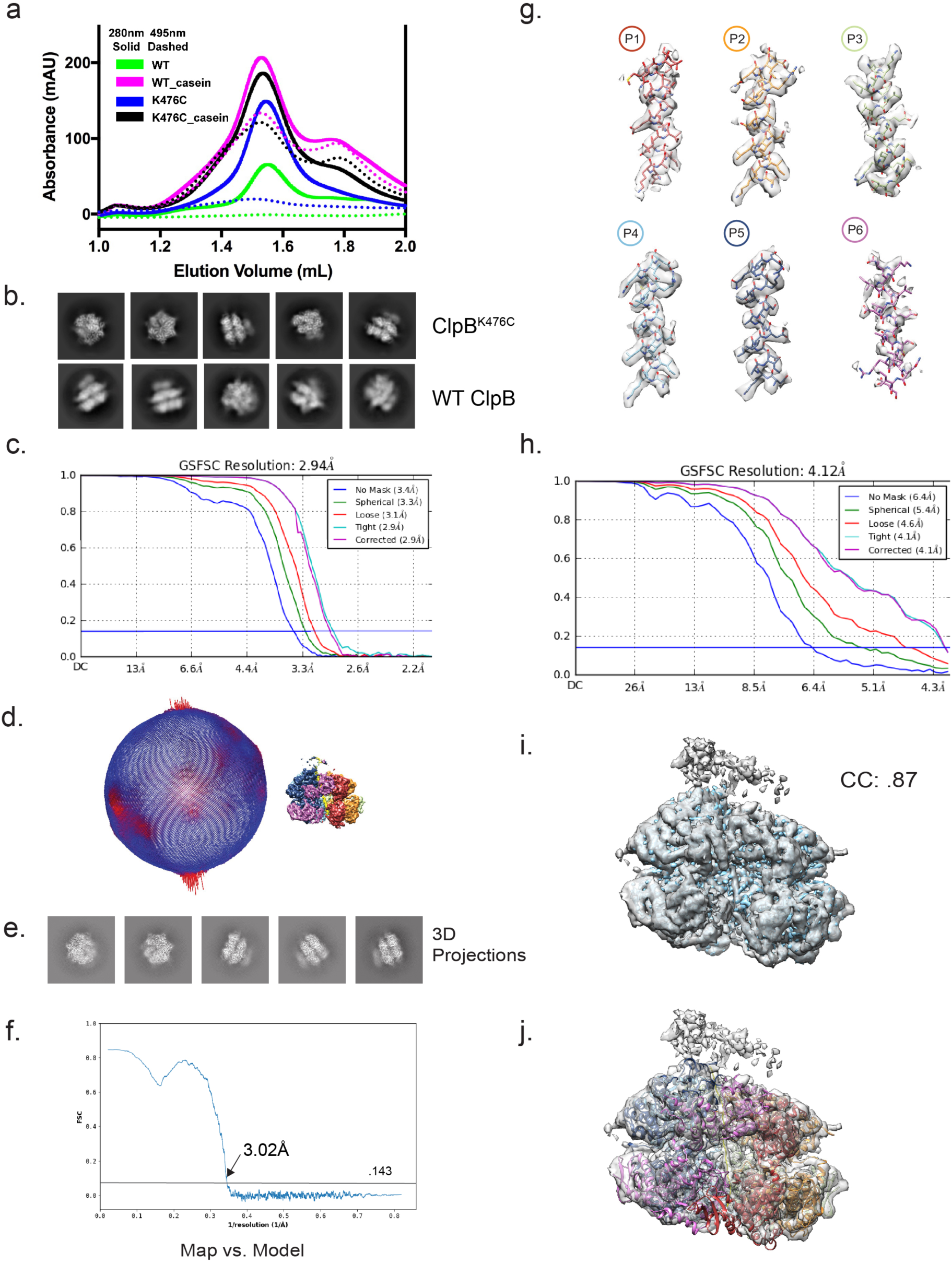
ATPγS-ClpB^K476C^:casein complex and analysis of cryo-EM data. **(a)** Formation of the substrate-bound WT ClpB and ClpB^K476C^ complexes by size exclusion chromatography (SEC) following incubation with FITC-casein and ATPγS. Absorbance traces are shown for λ = 280 (solid) for protein, and λ = 495 (dash) for FITC. **(b)** Reference-free 2D class averages of casein-bound WT and ClpB^K476C^ following SEC fractionation. **(c)** Gold standard FSC-curves of the final ClpB^K476C^:casein map using the total dataset following 2D classification. **(d)** Angular distribution of the particles for ClpB^K476C^:casein reconstruction determined using RELION^56^. **(e)** Projections of ClpB^K476C^:casein reconstruction showing top and side orientations that match reference-free averages in (b). **(f)** The map vs. model FSC-curve following atomic modeling in PHENIX^42^. **(g)** The cryo-EM map and atomic model of helix C3^7^ that includes residues 388–406 is shown for each protomer. **(h)** Gold standard FSC-curve of the final map of WT ClpB:casein. **(i)** Alignment of WT ClpB:casein (grey) and ClpB^K476C^:casein (blue) shown with the cross correlation (CC) value, calculated using UCSF Chimera^61^. **(j)** WT ClpB:casein map shown docked with the molecular model determined from the ClpB^K476C^:casein map.

**Supplementary Figure 2:**
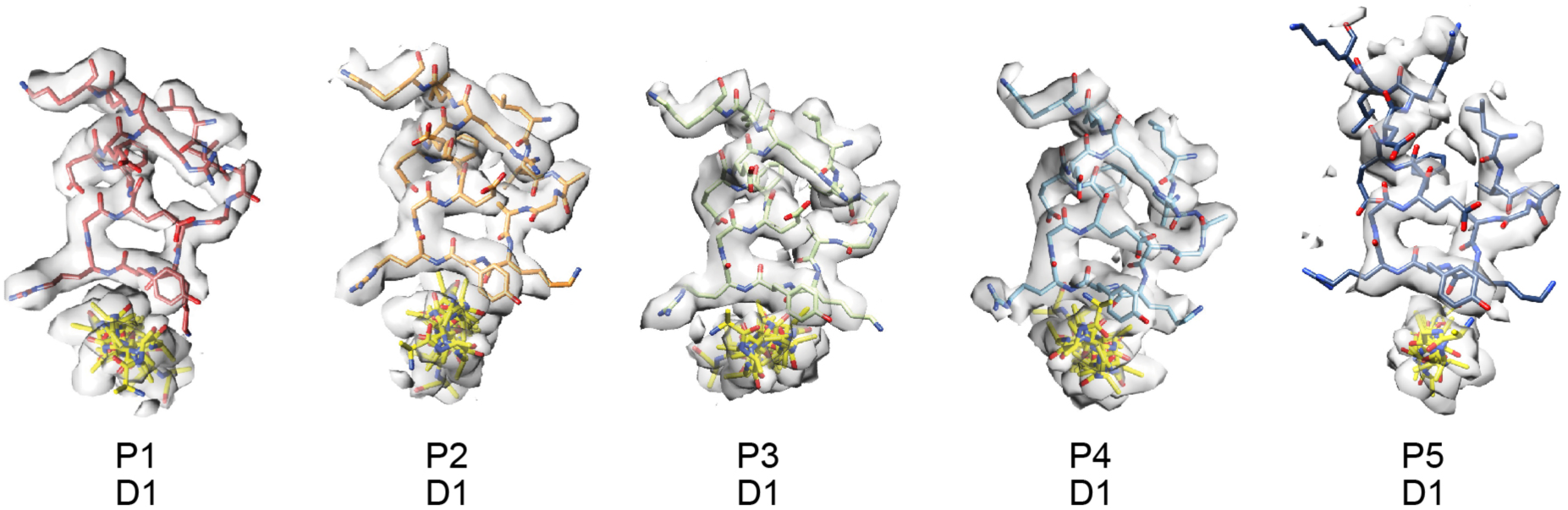
D1 pore loops have well-defined high-resolution structural features. Top view of the segmented map of the D1 loops from each protomer with residues (245–260) shown.

**Supplementary Fig. 3:**
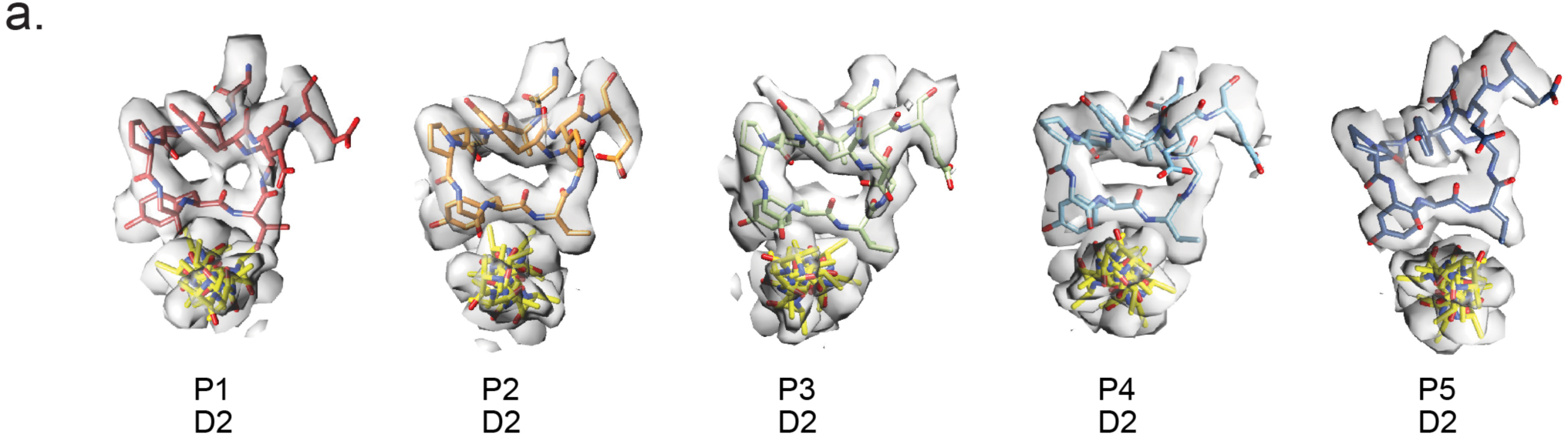
D2 pore loops have well-defined high-resolution structural features. Top view of the segmented map of the D2 loops from each protomer with residues (646–660) shown.

**Supplementary Figure 4:**
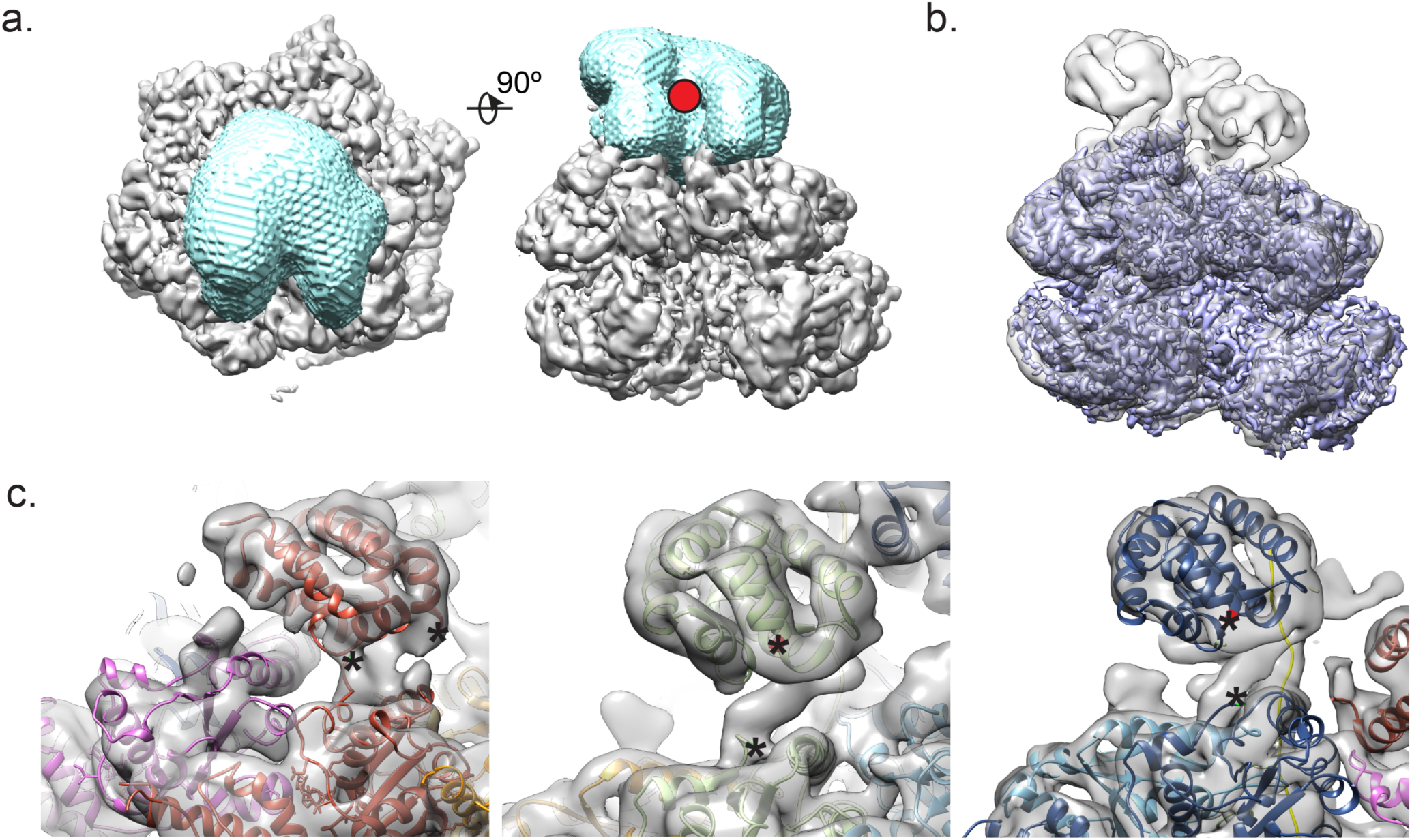
Extensive 3D and Focus classification improve resolution of NTDs. **(a)** Map and mask used in the focus classification of the NTDs. Red dot represents the new center of mass in which the particles were adjusted to. **(b)** The focus classification map after refinement (grey) is overlaid with the original map (purple). Both maps are shown at a threshold value of 4σ. **(c)** Map of the NTD refinement classification docked with the model, pdb: 1KHY, chain A with the full ClpB^K476C^:casein model. The C-terminus end of NBD1 (green), residue:161, and the N-terminus end of NTD (red), residue:142. To emphasize the ends, asterisks are also shown and are near the extra density that is present in which connection between NTD and NBD1 could be made.

**Supplementary Figure 5:**
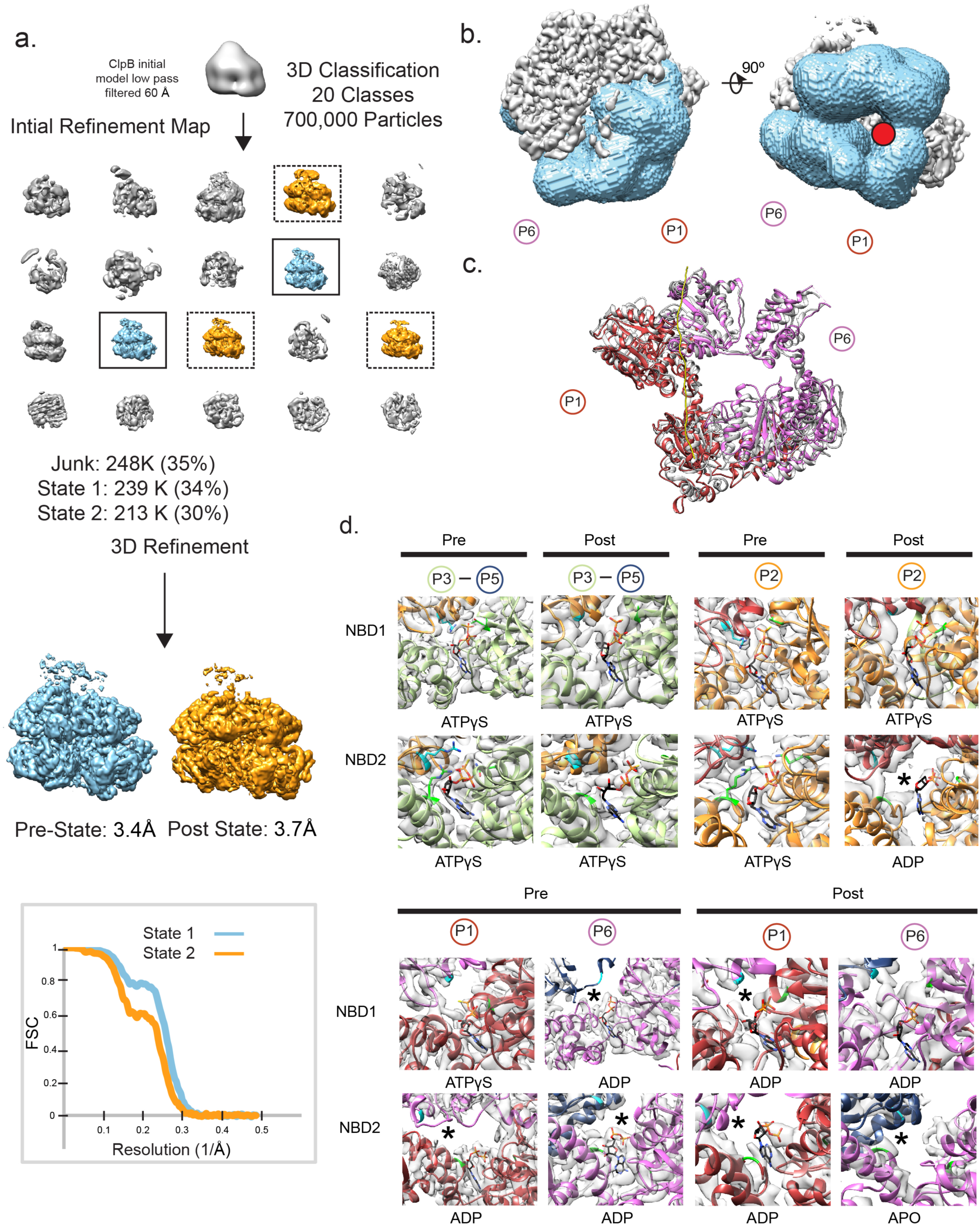
Extensive 3D and Focus classification identifies two different states of K476C-ClpB. **(a)** 3D classification scheme that was used for the ClpB^K476C^:casein dataset in identifying the two different states. The classes that were used in further refinement stages for each of the classes are identified by color, state 1 (blue) and state 2 (orange). Classes that were excluded from further refinement are indicated in grey. **(b)** Map and mask used in the focus classification of the two flexible front protomers, P1 and P6. Red dot represents the new center of mass in which the particles were adjusted to. **(c)** Model of pre-state (colored) aligned with 5ofo (grey), with only P1 and P6 shown. **(d)** Map and model shown to indicate nucleotide density within the nucleotide pockets of all six protomers, P1-P6, as well as both states, pre and post. The arg fingers (331 and 756) (cyan) and sensor residues (315,719,815) (green) are shown where density was present for them. Nucleotide pockets in which the arginine finger does not make contact with nucleotide is represented with asterisk.

**Supplementary Table 1:**
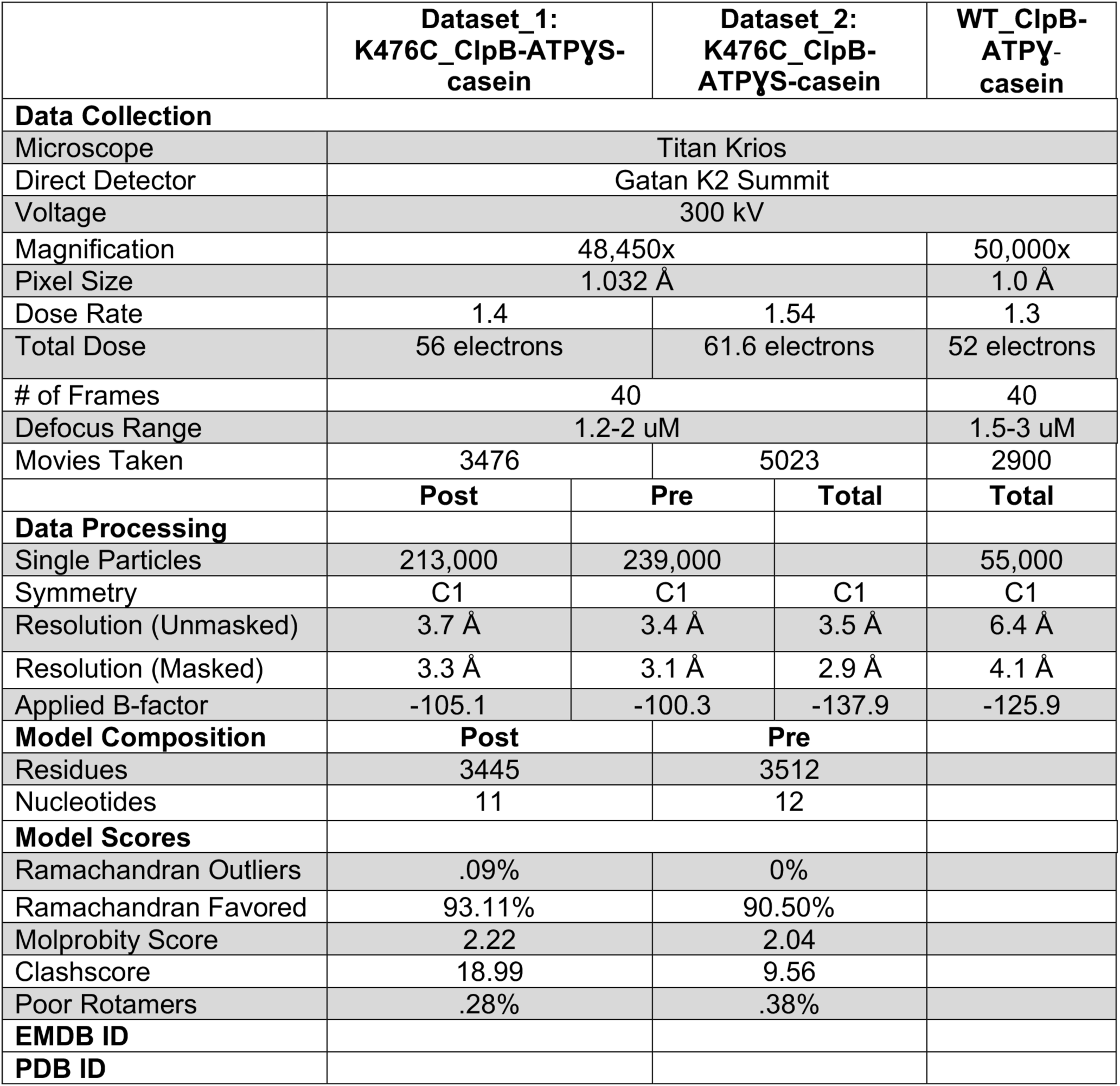
EM data collection and atomic model data. Information regarding data collection, processing, and atomic modeling parameters for the different structures.

**Supplementary Table 2:**
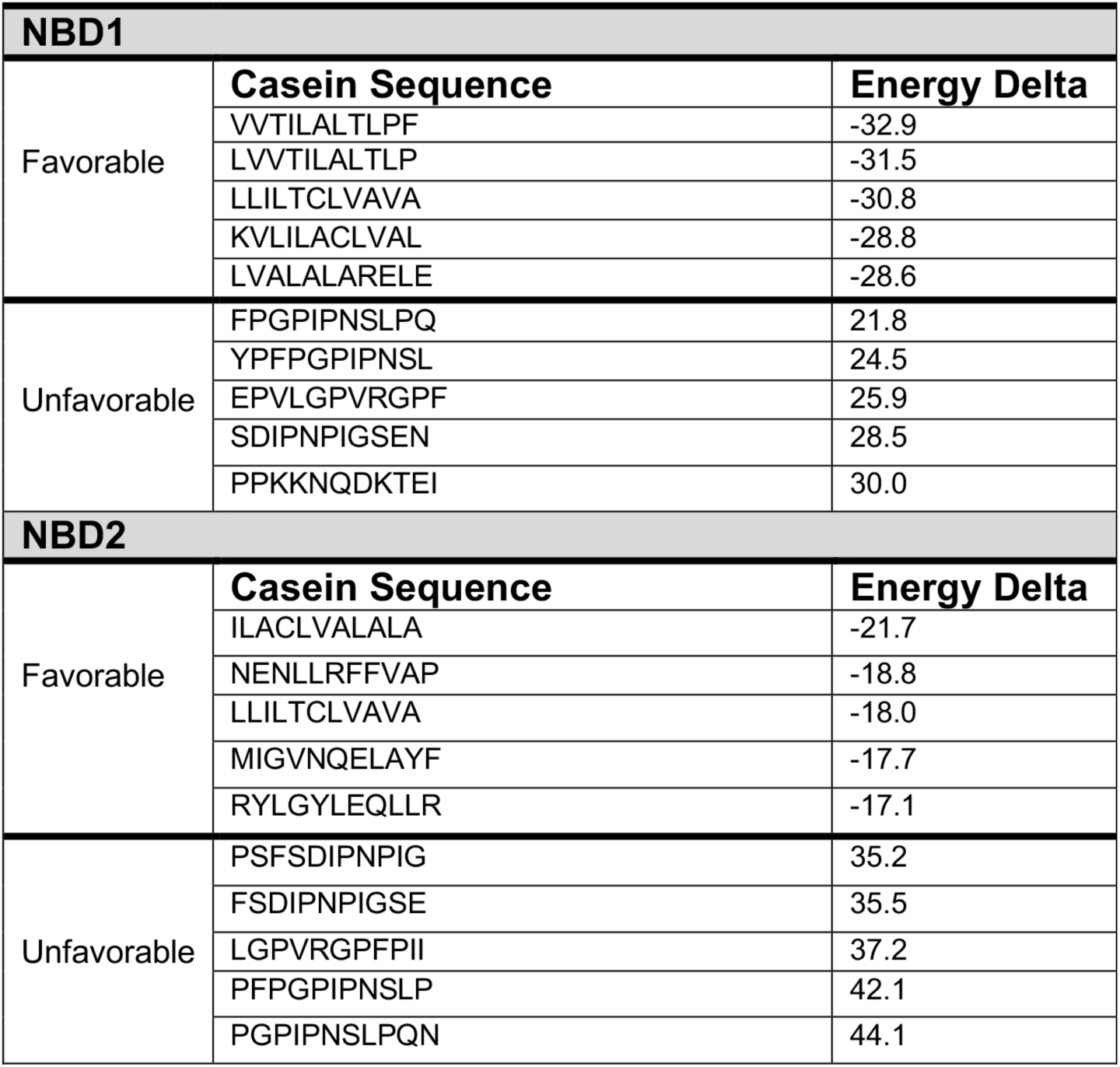
Casein Ssubstrate peptide modeling. Sequence and binding energy change for the five lowest (most favorable) and five highest (least favorable) scoring casein peptides.Movie showing a morph between the Pre and Post state conformations of the spiral seam protomers P1 and P6.

**Supplementary Movie 1: Pre and Post state conformational changes**. Movie showing a morph between the Pre and Post state conformations of the spiral seam protomers P1 and P6.

